# Comparative genomics of parasitoid lifestyle as exemplified by Mermithidae and Nematomorpha

**DOI:** 10.1101/2024.06.11.598342

**Authors:** Joseph Kirangwa, Viktoria Bednarski, Nadège Guiglielmoni, Robert Poulin, Eddy Dowle, Philipp H. Schiffer, Oleksandr Holovachov

## Abstract

Mermithidae and Nematomorpha are parasitoids united by the commonalities in their lifestyle – immature stages infect arthropod hosts, species from both phyla can manipulate their host to induce a similar water-seeking behaviour, and both have a final free-living non-feeding adult reproductive stage, often killing their host upon emergence. Some of these species are of great economic importance, being evaluated as biological control agents against mosquito vectors responsible for diseases like malaria, and other insect pests, but with scarce genomic resources currently available. Nematomorpha, despite being closely related to Nematoda, received insufficient attention in genomic research, leading to gaps in our understanding of their diverse genetic makeup. This study aimed to investigate the genetic features encoded in the genomes of both parasitoid taxa to identify similarities and parallels linked to their ecological lifestyles. We performed a comparative analysis of 12 genomes, comprising parasitoid, parasitic and free-living worms. The investigation revealed genomic signatures unique to parasitoid species, including expanded gene families enriched in neural transmission modulation, likely linked to the known host manipulation that both mermithids and nematomorphs exert on their hosts. The analysis also uncovered a diverse array of conserved transposable element superfamilies across both lineages. The findings from this study provide valuable insights into the potential genomic adaptations associated with parasitoidism in nematode and nematomorph worms. The identification of expanded gene families and conserved transposable element superfamilies sheds light on the molecular underpinnings of their unique biological traits. Additionally, the core set of orthologs specific to parasitoid worms offers new avenues for understanding the evolution of parasitism within these groups of organisms.

## Introduction

Mermithids (a family of the phylum Nematoda) and Nematomorphs (a phylum called horsehair worms or hairworms) represent two distinct groups of worms that exhibit intriguing resemblances in their parasitoid lifestyles and the alterations in behavior they induce in their definitive hosts. Parasitoids are a subgroup of parasites which “develop on or within another organism (=host), derive nourishment from it, and ultimately cause its demise during or at the end of this development” (modified after (Eggleton and Gaston 1990; Maggenti *et al*. 2005)). The parasitoid lifestyle is better known in insects Eggleton and Belshaw (1992) but there are important biological differences between them and parasitoid worms. In insects, the free-living adult feeds on a different source of nutrients and occupies a different ecological niche, compared to its parasitoid larvae. Thus, for parasitoid insects, killing the host imposes no cost on the immediate survival of individuals. The biology of parasitoid nematodes and nematomorphs is different. Adult mermithids and nematomorphs, as well as their invasive juveniles, are free-living and do not feed at all, overcoming the disadvantage of killing the host. The parasitoid juveniles of both mermithids and nematomorphs source their nutrition from a variety of arthropod species in a similar way (de Valdez 2005; Mishina *et al*. 2023; Sato *et al*. 2012). The cuticle of parasitoid juvenile nematodes undergoes molting within the host, and the epidermis endures a certain degree of degeneration (Schmidt-Rhaesa 2005), allowing for transcuticular feeding by the haemocoel-dwelling parasitoid (Rutherford *et al*. 1977; Skaling and MacKinnon 1988; Poinar Jr and Hess 1977), while its digestive system reduces in complexity (Schmidt-Rhaesa 2005). Accumulated nutrients are stored in the trophosome, a modified intestine, in Mermithidae and in parenchyma in Nematomorpha. This energy store allows free-living non-feeding adults to engage in mating and egg-laying for a long period of time, producing huge numbers of progeny, after emerging from the host (Hanelt *et al*. 2005; Nickle 1972).

Some Mermithidae and Nematomorpha are known to manipulate the behavior of their hosts by inducing water-seeking in terrestrial arthropods that harbour those worm species which complete their development in the aquatic environment (Maeyama *et al*. 1994; Vance 1996; Poulin 1998; Thomas *et al*. 2002; Ponton *et al*. 2011; Allahverdipour *et al*. 2019; Herbison *et al*. 2019b). In nematomorphs, this manipulation may involve alterations in neurotransmitter levels and the regulation of the host’s nervous system through parasite-produced molecules like serotonin, dopamine, and octopamine (in *Paragordius tricuspidatus*) (Thomas *et al*. 2003) and Wnt family proteins (e.g. in *Paragordius tricuspidatus*, and *Spinochor-dodes tellinii*) Biron *et al*. (2005, 2006). In mermithids, the species *Thaumamermis zealandica* induces an increase in haemolymph osmolality in its semi-terrestrial crustacean host *Talorchestia quoyana* – this alteration may create a requirement for water in the host, providing a potential explanation for the observed water-seeking behavior (Williams *et al*. 2004). Another mermithid species, *Mermis nigrescens*, induces erratic or hyperactive behavior in the European earwig *Forficula auricularia*, potentially through initial modulation of Mtp*α* (Mitochondrial trifunctional protein *α* subunit) and subsequent manipulation of neuronal connections and neurotransmitters, ultimately leading to a hydrophilic state. Clathrin is a key protein in this manipulation process (Herbison *et al*. 2019a). Regulation in host brains, coupled with predominant proteomic changes in hosts infected by large worms, and shared functional characteristics of proteins regulated in mermithid and hairworm-infected hosts, especially in axon or dendrite and synapse modulation, strongly support the notion of adaptive manipulation of host behavior.

Substantial differences between mermithids and nematomorphs relate to their free-living stages. The infective juveniles of mermithids are actively seeking their host and use digestive enzymes to penetrate the body wall and make their way to the haemocoel (Shamseldean and Platzer 1989), whereas the larvae of nematomorphs are semi-sessile and rely on the intermediate paratenic host to reach the definitive host (Hanelt *et al*. 2005). Mermithid adults are armed with a variety of sensory structures and likely engage them in host-finding (Robinson *et al*. 1990) and mate-finding (Burr and Babinszki 1990; Burr *et al*. 2000). Our knowledge of the receptors present in Nematomorpha is limited – existing ultrastructural studies provide minimal insights into their functionality and role (Schmidt-Rhaesa 2004; Sokolova *et al*. 2022). From the phylogenetic point of view, the adoption of a parasitoid lifestyle in all extant species of Nematomorpha suggests that this characteristic likely originated early in the evolutionary history of the group, and is synapomorphic to the phylum. Within the phylum Nematoda, a parasitoid lifestyle is not unique and originated independently in four different lineages (i.e. is homoplasic): Mermithidae, Marimermithidae, Benthimermithidae and *Neocamacolaimus* (Miljutin 2013; Tchesunov *et al*. 2023; Holovachov and Boström 2014) but only Mermithidae were evolutionary successful, with about one thousand known species infesting a broad variety of hosts.

Given the significant representation of parasitic organisms among diverse species, their emergence across various phyla underscores a vast biological spectrum Poulin (2011). Notably, taxa such as mermithids and nematormorphs exhibit convergence in functionality and phenotype. We investigate how this convergence is demonstrated at the genomic level. Genomic resources for both Nematomorphs and Mermithids are still limited, but this paper introduces a new genome for *Romanomermis iyengari*. The sizes of few published genomes range from 250-500 Mb and the genomes overall exhibit a reduced gene content when compared to metazoan orthologs. (Cunha *et al*. 2023; Eleftheriadi *et al*. 2023; Guiglielmoni *et al*. 2024; Bhattarai *et al*. 2024; Kirangwa *et al*. 2023). A comprehensive comparative analysis of these genomes within an evolutionary framework remains an unexplored aspect. Here we provide the first comparative analysis to gain an understanding of their genome structure in general, and the likely evolutionary drivers and mechanisms behind their specialized parasitoid lifestyle in particular. We focus specifically on those possible genetic factors that are crucial for parasite survival and those involved in the interaction with their definitive hosts. We conducted inter-phyla comparative analyses of gene families, focusing on expansion vs. contraction and gain vs. loss, looking for common patterns during the evolution of these two distantly related but ecologically convergent parasitoid lineages. Furthermore, we investigate the contribution of transposable elements (TE) to genome complexity. Mobile TEs often induce deleterious effects upon insertion through inactivation of genes but infrequently also contribute to genome dynamics and adaptability by reorganization and introduction of novelties (Xiaoqian Jiang 2017). They were found to be very abundant in the genome of an insect parasitoid species *Nasonia* (Wurm and Keller 2010), although the significance of this remains unclear. Thus, we also analyzed the transposable elements in worm parasitoids to elucidate their prevalence and distribution, paving the way towards understanding their role in evolution of the parasitoid lifestyle.

## Materials and methods

### Data Collection

We conducted a comparative genomics analysis encompassing all published and one new genome of parasitoid worms, including three mermithid species (*Romanomermis culicivorax, Romanomermis iyengari, Mermis nigrescens*), and four nematomorph species (*Acutogordius australiensis, Gordionus montsenyensis, Gordius aquaticus*, and *Nectonema munidae*) (Guiglielmoni *et al*. 2024; Bhattarai *et al*. 2024; Eleftheriadi *et al*. 2023; Cunha *et al*. 2023). Additionally, our analysis includes closely related in-group nematode parasites, such as *Trichinella spiralis, Onchocerca volvulus*, and *Brugia malayi*, as well as free-living limno-terrestrial species *Plectus sambesii* and *Mesodorylaimus* sp. Supplementary material (Table 1) offers further details on the genomic characteristics of these species and their respective sources. All subsequent data analyses and interpretations were conducted within a phylogenetic context, utilizing published tree topologies as a reference (Ahmed *et al*. 2022; Cunha *et al*. 2023).

**Table 1.** Top 20 GO functional terms enriched in Mermithid gene family orthologous clusters.

| GO ID | Description | p value |
| --- | --- | --- |
| GO:0007165 | signal transduction | 1.02E-69 |
| GO:0006511 | ubiquitin-dependent protein catabolic process | 2.85E-53 |
| GO:0016192 | vesicle-mediated transport | 2.40E-37 |
| GO:0015031 | protein transport | 6.25E-33 |
| GO:0007283 | spermatogenesis | 1.81E-23 |
| GO:0007601 | visual perception | 1.63E-22 |
| GO:0046872 | metal ion binding | 2.41E-21 |
| GO:0008270 | zinc ion binding | 1.80E-20 |
| GO:0006508 | proteolysis | 3.63E-19 |
| GO:0016567 | protein ubiquitination | 5.38E-17 |
| GO:0006099 | tricarboxylic acid cycle | 1.13E-13 |
| GO:0006468 | protein phosphorylation | 3.3E-12 |
| GO:0006814 | sodium ion transport | 9.83E-11 |
| GO:0034765 | regulation of ion transmembrane transport | 2.39E-8 |
| GO:0005634 | nucleus | 4.61E-8 |
| GO:0097225 | sperm midpiece | 4.62E-8 |
| GO:0004867 | serine-type endopeptidase inhibitor activity | 1.05E-7 |
| GO:0007224 | smoothened signaling pathway | 5.74E-7 |
| GO:0046676 | negative regulation of insulin secretion | 4.67E-7 |
| GO:0034976 | response to endoplasmic reticulum stress | 1.83E-5 |

### HMW DNA extraction

The *Romanomermis iyengari* nematodes were selected from moss samples provided by Professor Dr. Edward Platzer at the University of California Riverside. High-molecular-weight DNA from 20 individuals of *Romanomermis iyengari* was extracted using a salting out protocol as follows. The nematodes were collected and washed in water, then flash-frozen using liquid nitrogen in a salt-based extraction buffer (Tris-HCl 100 mM, ethylenediaminetetraacetic acid 50 mM, NaCl 0.5 M and sodium dodecylsulfate 1 %). Samples were incubated overnight at 55°C after addition of 5 µL of proteinase K. DNA was precipitated using NaCl 5 M, yeast tRNA and isopropanol, and incubated at room temperature for 30 minutes, then pelleted at 18,000 g for 20 minutes (4°C). The DNA was washed twice with 80 % ethanol and spun at 18,000 g for 10 min (4°C). The DNA pellet was eluted in an elution buffer (D3004-4-10 Zymo Research) and incubated at 50°C for 10 minutes. RNA was removed by incubating with RNAse (Qiagen, 19101) for 1 hour at (37°C). DNA concentrations were quantified using a Qubit 4 fluoremeter with 1X dsDNA kit. HiFi libraries were prepared with the Express 2.0 Template kit (Pacific Biosciences, Menlo Park, CA, USA) and sequenced on a Sequel II/Sequel IIe instrument with 30 hours movie time. HiFi reads were generated using SMRT Link (v10,Pacific Biosciences, Menlo Park, CA, USA) with default parameters.

### Genome assembly and annotation of *Romanomermis iyengari*

PacBio HiFi reads were assembled using NextDenovo v2.5.2 (Hu *et al*. 2023). Assembly statistics were generated using assembly-stats v1.0.1, and ortholog completeness was assessed using the Benchmarking Universal Single-Copy Orthologs (BUSCO) tool v5.4.7 with parameters -m genome against the metazoa_odb10 and nematoda_odb10 lineages. Subsequently, PacBio HiFi reads were aligned to the HiFi assembly using minimap2 v2.24 with parameters -ax map-hifi, and the resulting mapped reads were sorted using SAMtools v1.6. Contigs were aligned against the nt database using BLAST v2.13.0, and the outputs were processed using BlobToolKit v4.1.5 to remove contaminants. Reads were remapped using minimap2 v2.24, and the output was provided to purge dups v1.2.5 to eliminate uncollapsed haplotypes and assembly statistics were recalculated using assembly-stats v1.0.1. Protein-coding genes were predicted using the BRAKER3 pipeline (Gabriel *et al*. 2023), leveraging protein alignments from related species as evidence. To construct a protein database, proteins from *Romanomermis culicivorax* and *Mermis nigrescens* were combined with OrthoDB (Kuznetsov *et al*. 2023) v11 Metazoa and Eukaryota gene orthologs. The protein database was used as input to BRAKER3 for gene prediction in the soft-masked genome of *Romanomermis iyengari*. Subsequently, BUSCO analyses were conducted (utilizing the nematoda_odb10 and metazoa_odb10 lineages) on the predicted protein sequences to evaluate biological completeness. Additionally, each gene set was compared with the proteomes from other species included in this study by clustering the longest isoform of each predicted gene using OrthoFinder (Emms and Kelly 2019) v2.5.5.

### Orthologous cluster identification and visualization

Orthologous gene identification was conducted using OrthoFinder (Emms and Kelly 2019) integrated into OrthoVenn3 (Sun *et al*. 2023), employing an E-value cutoff of 1e-5 for comparing all protein similarities and an inflation value of 1.5 to generate orthologous clusters via the Markov clustering algorithm. Gene Ontology term annotations for individual orthologous clusters were further performed.

### Gene family analysis

In order to gain insight into the evolutionary dynamics of genes within both Mermithidae and Nematomorpha, we assessed the expansion and contraction of gene ortholog clusters. Computational analysis of gene family evolution was conducted using CAFE5 (Mendes *et al*. 2020) software, defining expansion and contraction based on differences in cluster size between ancestors and each current species. Furthermore, to elucidate the genetic factors or mechanisms underlying phenotypic traits in species, the gene families were annotated with GO terms.

### Repeat sequence analysis

To assess the content of transposable elements (TE) within the Mermithidae and Nematomorpha, first, a TE and repeat library was built using EDTA v2.1.3 (Ou *et al*. 2019) with parameters **– sensitive 1 –anno 1**. TEs were then annotated and classified to superfamily level using the FasTE pipeline (Ellen A. Bell 2022). Output files were parsed with the R script “RMTrips”, contained in FasTE. A custom R script “RMTrips output to divsum-file format” was used, renaming the TE superfamilies according to Wicker-classification (Wicker *et al*. 2007), and converting the data format to further plot the TE divergence landscapes with the “Plot Kimura Distance” R script. Stacked barplots were plotted in R.

## Results

### The genome sequence of *Romanomermis iyengari*

The assembly size of *Romanomermis iyengari* genome is 304.4 Mb, featuring a 37 % GC content and scaffold N50 of 326 kb. The longest scaffold stretching over 3.9 Mb. Gene annotation prediction yielded 16,503 protein coding genes. Overall BUSCO scores are 66.8 % completeness for Metazoa, with duplicated orthologs at 13.2 % and fragmented orthologs at 3.4 %. In terms of Nematoda, BUSCO assessment shows 50.4 % completeness, with duplicated orthologs at 13.1 % and fragmented orthologs at 1.9 %. Moreover, 54.28 % of the *R. iyengari* genome has been identified as repetitive elements.

### Identification and analysis of orthologous clusters in Mermithidae and Nematomorpha

In our analysis, a set of 335 highly conserved orthologous clusters were unique to mermithids and 263 to Nematomorpha respectively (Figure 1).

**Figure 1.**
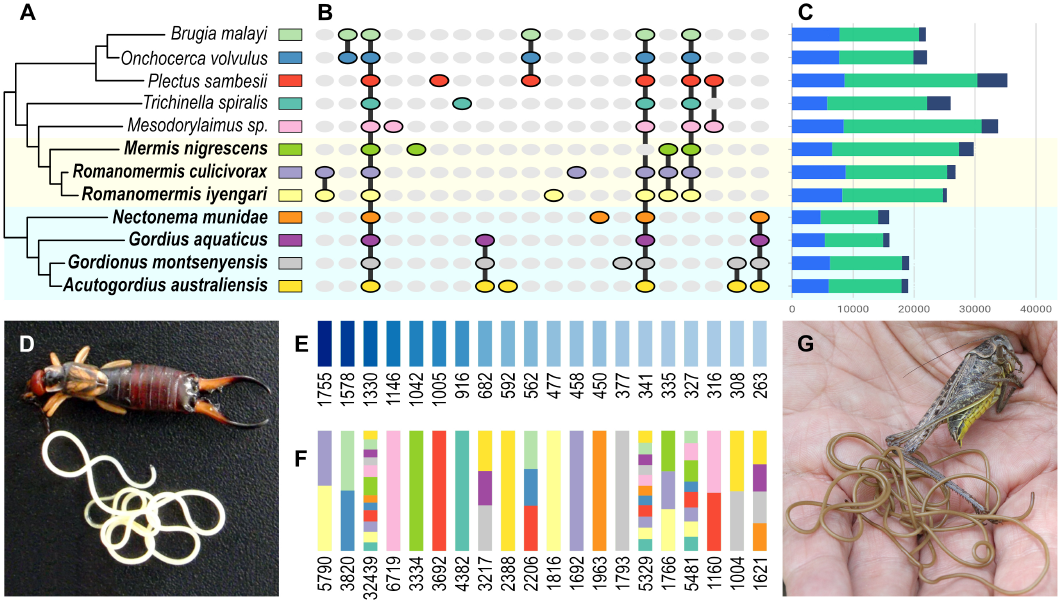
Orthologous cluster analysis: **(A)** Phylogenetic relationships between studied species (color label corresponds with colors in B and F); **(B)** Distribution of shared and unique orthologous gene clusters among studied species, colors correspond to species in A; **(C)** Number of protein sequences (blue), orthologous clusters (green) and singletons (dark purple); **(D)** Mermithid *Mermis nigrescens* emerging from its host (photograph by Bronwen Presswell, reproduced with permission); **(E)** Number of shared orthologous clusters for each subset of species or the number of unique clusters; **(F)** Combined number of proteins in shared and unique clusters, colored segments represent relative proportion of proteins for each species, colors correspond to species in A; **(G)** Nematomorph emerging from its host (photograph by Alastar Rae, reproduced here under CC BY-SA 2.0 license).

Mermithid orthologous clusters were over-represented for 40 Gene Ontology (GO) terms. Here, the top 20 enriched GO terms based on `e-value` *<* 1 *×* 10^−5^ are presented (Table 1). However, additional enriched functional signatures were observed, including regulation of DNA methylation (GO:0044030), G-protein coupled receptor activity (GO:0004930), and chloride channel activity (GO:0005254). A comprehensive list of all enriched GO terms is provided as supplementary material (File_S2_Mermithids_orthologs).

We found no orthologous clusters that would be uniquely shared by mermithids and nematomorphs and at the same time completely absent in other species included in the analysis.

Nematomorpha orthologous gene clusters were over-represented for 76 Gene Ontology (GO) terms. The top 20 enriched orthologous clusters were enriched in GO terms are shown in (Table 2). Other enriched orthologous clusters beyond the top 20 were enriched in sodium ion transport (GO:0006814), response to toxic substance (GO:0009636), response to starvation (GO:0042594), and sensory perception of sound (GO:0007605). A complete list is shown in the supplementary material (File_S3_Nematomorphs_orthologs).

**Table 2.** Top 20 GO functional terms enriched in Nematomorph gene family orthologous clusters.

| GO ID | Description | p value |
| --- | --- | --- |
| GO:0007165 | signal transduction | 5.72E-142 |
| GO:0006511 | ubiquitin-dependent protein catabolic process | 5.19E-87 |
| GO:0016192 | vesicle-mediated transport | 1.11E-62 |
| GO:0016021 | integral component of membrane | 7.67E-62 |
| GO:0006355 | regulation of transcription, DNA-templated | 2.54E-59 |
| GO:0015031 | protein transport | 4.36E-56 |
| GO:0007275 | multicellular organismal development | 2.10E-49 |
| GO:0008270 | zinc ion binding | 3.45E-47 |
| GO:0006508 | proteolysis | 1.10E-46 |
| GO:0016032 | viral process | 6.19E-41 |
| GO:0016055 | Wnt signaling pathway | 2.62E-39 |
| GO:0006468 | protein phosphorylation | 7.69E-36 |
| GO:0006357 | regulation of transcription from RNA polymerase II promoter | 2.89E-34 |
| GO:0007601 | visual perception | 7.13E-34 |
| GO:0016567 | protein ubiquitination | 4.72E-26 |
| GO:0006486 | protein glycosylation | 1.38E-21 |
| GO:0030433 | ER-associated ubiquitin-dependent protein catabolic process | 2.45E-20 |
| GO:0006886 | intracellular protein transport | 5.58E-17 |
| GO:0071805 | potassium ion transmembrane transport | 7.80E-17 |
| GO:0034765 | regulation of ion transmembrane transport | 3.67E-16 |

### Gene family evolution in Mermithidae and Nematomorpha

In our investigation of Mermithidae we identified 226 expanded gene families and 114 contracted gene families in *Mermis nigrescens*, 89 expanded gene families and 122 contracted gene families in *Romanomermis culicivorax* and 96 expanded gene families and 145 contracted gene families in *Romanomermis iyengari* (Figure 2). Within the lineage leading to the Mermithidae family, we identified 169 gene families that underwent contraction and 40 gene families that experienced expansion.

**Figure 2.**
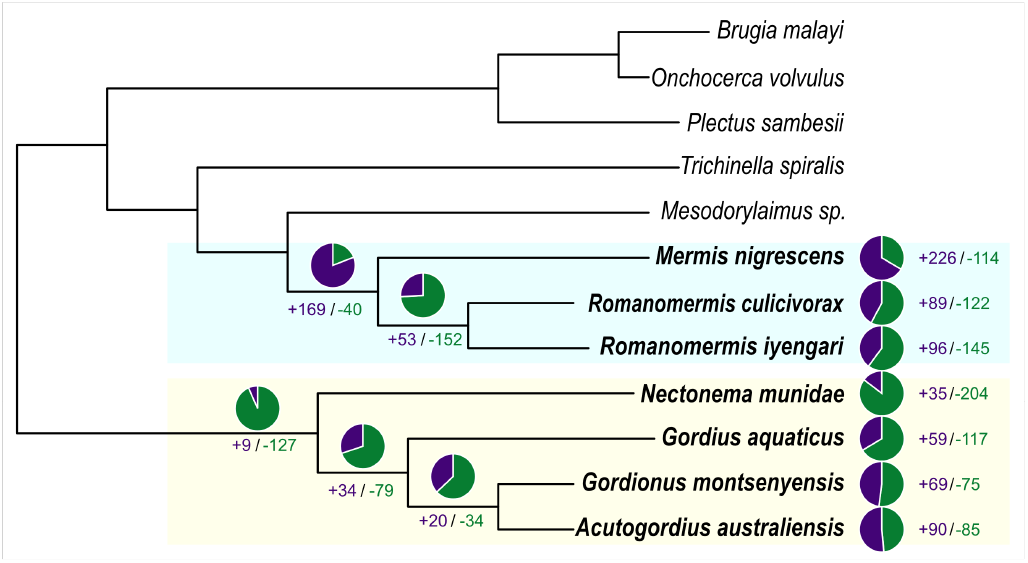
Pie charts show gene family size. The number of contracted (purple) and expanded (green) gene families. The Cladogram on the left shows the evolutionary history of gene families and differences between species included in the analysis.

Table 3 displays the functional annotation with GO terms of the expanded gene families predominantly enriched in mermithid nematodes. Though the top 20 are shown, mermithids exhibited enrichment in various other signatures, including norepinephrine transport (GO:0015874), motor activity (GO:0003774), neurotransmitter transport (GO:0006836), response to toxic substance (GO:0009636), response to oxidative stress (GO:0006979), regulation of synapse structural plasticity (GO:0051823), telomere maintenance via recombination (GO:0000722), response to starvation (GO:0042594), spermatogenesis (GO:0007283), DNA recombination and GABA-A receptor activity (GO:0004890). They are shown in the supplementary material (File_S4_mermithid_expanded_gene_families). The top five contracted gene families were enriched in the structural constituent of the cuticle (GO:0042302), protein heterodimerization activity (GO:0046982), nucleosome assembly (GO:0006334), macromolecular complex binding (GO:0044877), peptidyl-proline hydroxylation to 4-hydroxy-L-proline (GO:0018401). A comprehensive list of all contracted gene families is given in the supplementary material (File_S6_mermithid_contracted_gene_families).

**Table 3.** Top 20 GO functional terms enriched in Mermithid expanded gene family clusters.

| GO ID | Description | p value |
| --- | --- | --- |
| GO:0019346 | transsulfuration | 9.73E-82 |
| GO:00304311 | sleep | 1.45E-56 |
| GO:0006412 | translation | 2.20E-58 |
| GO:0006511 | ubiquitin-dependent protein catabolic process | 3.21E-51 |
| GO:0015267 | channel activity | 4.63E-49 |
| GO:0005198 | oligosaccharide metabolic process | 3.99E-47 |
| GO:0005198 | structural molecule activity | 3.84E-43 |
| GO:0005993 | trehalose catabolic process | 1.31E-42 |
| GO:0005991 | trehalose metabolic process | 1.27E-40 |
| GO:0007017 | microtubule-based process | 3.41E-40 |
| GO:0016185 | synaptic vesicle budding from presynaptic membrane | 4.59E-38 |
| GO:0016021 | integral component of membrane | 1.74E-36 |
| GO:0015144 | carbohydrate transmembrane transporter activity | 1.02E-32 |
| GO:0018107 | peptidyl-threonine phosphorylation | 2.84E-32 |
| GO:0006556 | S-adenosylmethionine biosynthetic process | 6.67E-32 |
| GO:0001505 | regulation of neurotransmitter levels | 9.84E-32 |
| GO:0006833 | water transport | 6.67E-31 |
| GO:0003774 | motor activity | 5.77E-29 |
| GO:0007016 | cytoskeletal anchoring at plasma membrane | 8.69E-29 |
| GO:0045332 | phospholipid translocation | 8.72E-29 |

We identified 90 expanded gene families and 85 contracted gene families in *Acutogordius australiensis*, 69 expanded gene families and 75 contracted gene families in *Gordionus montsenyensis*, 59 expanded gene families and 117 contracted gene families in *Gordius aquaticus*, and 35 expanded gene families and 204 contracted gene families in *Nectonema munidae*. We found 9 expanded gene families and 127 contracted gene families in the MRCA of the analysed nematomorph species (Figure 2). In nematomorphs, the expanded gene families were most enriched in GO terms shown in Table 4. Additionally, several others were significantly enriched in for example norepinephrine transport (GO:0015874) and neurotransmitter transport (GO:0006836), along with DNA-mediated transposition (GO:0006313), spermatogenesis (GO:0007283). Further details on these findings can be found in the supplemen-tary material (File_S5_nematomorphs_expanded_gene_families). The top five contracted gene families based on e-value *<* 1 *×* 10^−5^ were enriched in structural constituent of the cu-ticle (GO:0042302), microtubule-based process (GO:0007017), translation (GO:0006412), channel activity (GO:0015267) and oligosaccharide metabolic process (GO:0009311). A full list of all contracted gene families is in supplementary material (File_S7_nematomorphs_contracted_gene_families).

**Table 4.**
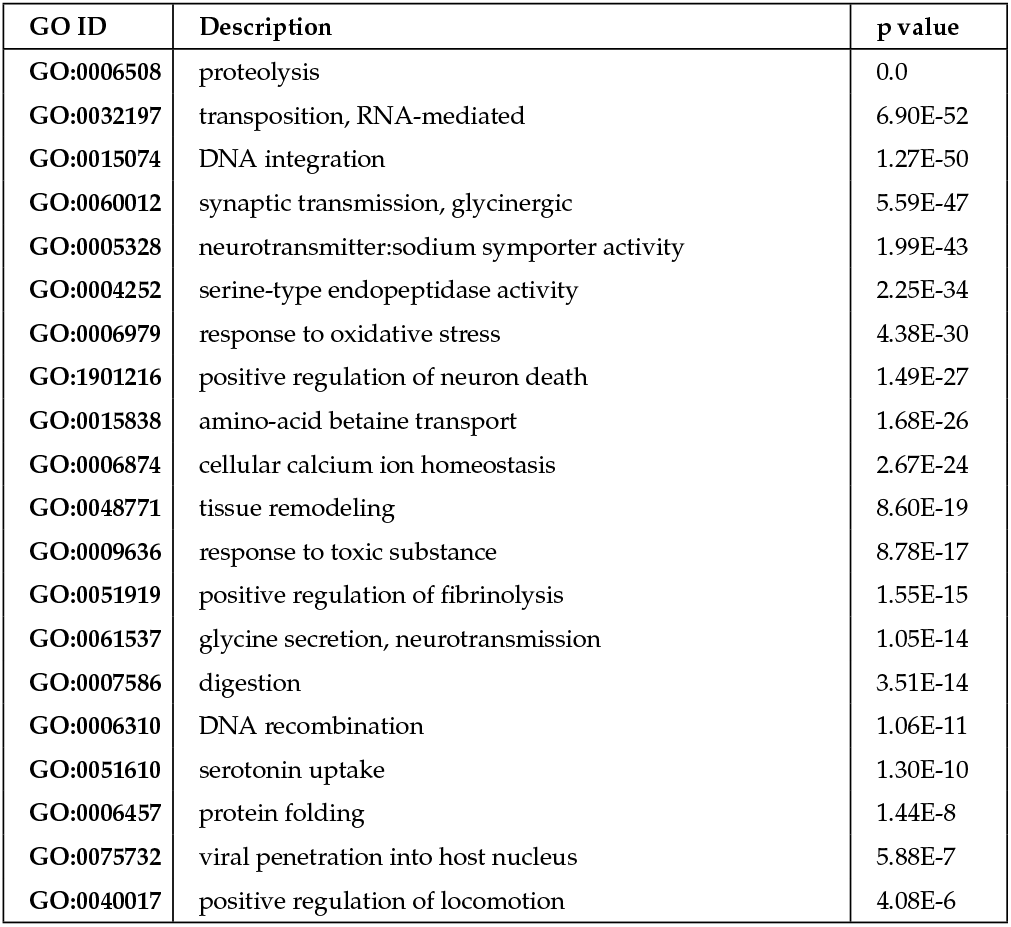
Top 20 GO functional terms enriched in Nematomorph expanded gene family clusters.

### Transposable elements

TE and repeat divergence landscapes are useful to infer recent and past activity of TEs, as TEs accumulate mutations and degrade over time. The Kimura substitution level is a measure of divergence from the TE consensus sequence. Highly diverged sequences are likely to have lost the ability to transpose and become inactive remnants. Both nematomorph and mermithid genomes exhibit a high proportion of transposable elements. They display a history of high TE activity and even recent bursts. In our analysis, *Nectonema munidae* contains TEs at various divergence levels. Prominent TE superfamilies include DTA_hAT and DTA_hAT_MITE, showing significant recent and ancient activity. *Gordius aquaticus* shows a broad distribution of TEs with peaks at low and intermediate divergence levels. The profiles of *Gordionus montsenyensis* and *Acutogordius australiensis* display a high percentage of TEs, with significant activity at multiple divergence levels. *Mermis nigrescens* shows significant TE presence with peaks at various divergence levels while *Romanomermis culicivorax* shows multiple peaks, similar to other mermithids. *Romanomermis iyengari* displays a high percentage of TEs, with significant activity at multiple divergence levels. *Mesodorylaimus sp*. shows low overall TE content with a few peaks, suggesting limited recent TE activity. *Plectus sambesii* and *Trichinella spiralis* reveal striking recent activity, while *Brugia malayi* shows intermediate increases in recent activity. *Onchocerca volvulus* displays low TE activity. (Figure 3).

**Figure 3.**
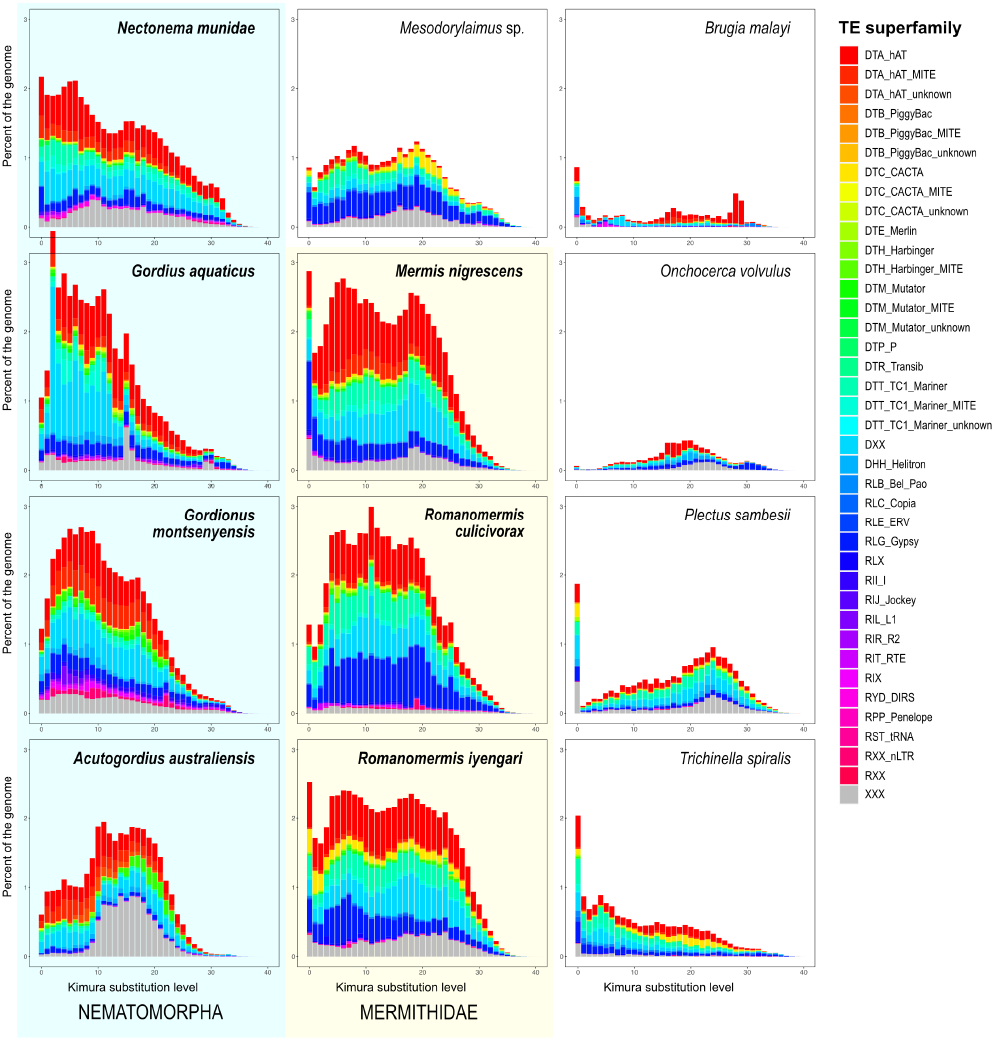
TE and repeat divergence landscapes showing TE portions as percent of the genome (y-axis) over Kimura substitution levels (x-axis) obtained via the “FasTE” pipeline. Legend for TE superfamily colors is shown to the right. XXX corresponds to unclassified repeats.

All studied species of Nematomorpha have a comparable genome size between 200-300 Mb. Nematomorpha and Mermithidae species generally exhibit a high combined length of TEs within their genomes, indicating extensive TE content. Other Nematodes show a lower combined length of TEs (Figure 5). *Mermis nigrescens* displays the highest combined TE length among the Mermithidae. *Brugia malayi* and *Onchocerca volvulus* Show relatively low TE content. (Figure 4). *Plectus sambesii* and *Trichinella spiralis* reveal slightly higher TE content than *Brugia malayi* and *Onchocerca volvulus*, but still much lower than Nematomorpha and Mermithidae species. *Mesodorylaimus sp*. has moderate TE content, with contributions from multiple superfamilies. Comparing the composition of TEs it becomes evident that DNA elements of the superfamilies *hAT, CACTA, Mutator, Transib* and *Mariner* are conserved between the species, while also containing a large portion of not further discriminated DNA elements. Further, all genomes contain Helitron elements and the composition of retroelements is dominated by the superfamily *Gypsy* (Figure 5).

**Figure 4.**
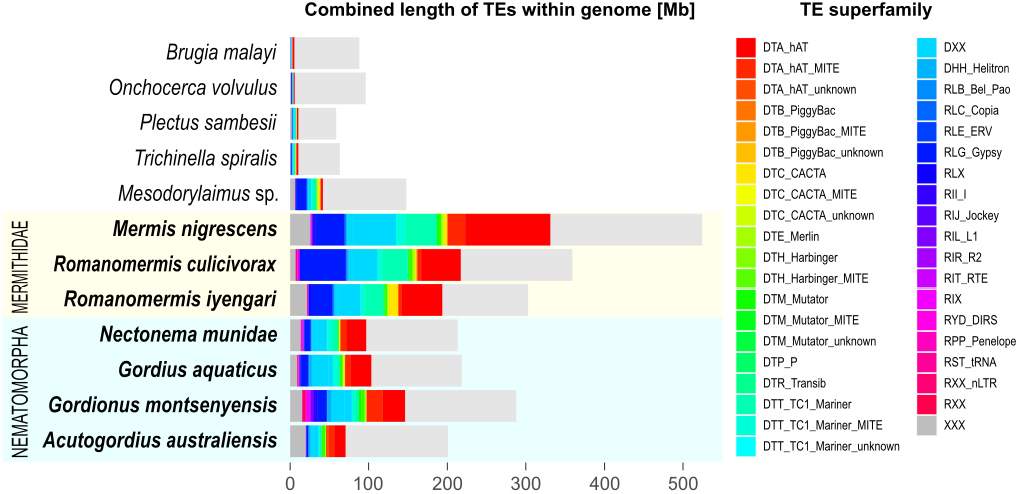
Stacked barplots show the amounts of TEs in Mb within the genomes of species included in the analysis.

**Figure 5.**
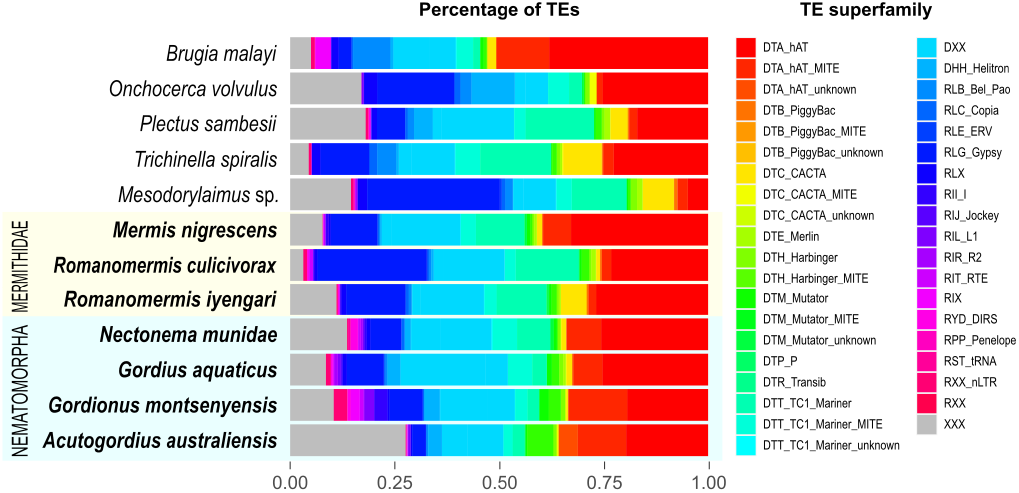
Stacked barplots show relative amounts of TEs within the genomes of species included in the analysis.

## Discussion

### The genome of *Romanomermis iyengari*

*Mermis nigrescens* and *R. culicivorax* stand out as the only mermithid nematodes with publicly available whole-genome assemblies to date Bhattarai *et al*. (2024); Guiglielmoni *et al*. (2024). These genomes exhibit assembly sizes ranging from 300 to 524 Mb, boasting N50’s exceeding 1 Mb. Notably, these assemblies surpass the completeness and contiguity of the *R. iyengari* genome presented in our current study which was generated with only PacBio HiFi long read data. However, it is noteworthy that the repeat content is high in all these three genomes (50-84 %). While the exact implications of this elevated repeat content remain unclear, further investigations may unveil its biological significance. These newly generated resources and insights promise to enhance our understanding of the genomic architecture and biology of mermithid nematodes, shedding light on their adaptation to a diverse array of insect hosts.

### Gene family evolution in Mermithidae and Nematomorpha

We identified significant expansions and contractions in gene families across all analyzed species of the family Mermithidae (*Mer-mis nigrescens, Romanomermis culicivorax*, and *Romanomermis iyengari*). The analysis of nematomorph species revealed varying patterns of expansions and contractions in gene families in studied species (*Acutogordius australiensis, Gordionus montsenyensis, Gordius aquaticus*, and *Nectonema munidae*) but at a lesser scale than in mermithids. Both combined, however, provide interesting insight into their evolutionary genomic dynamics in general and point towards gene families that may be associated with their parasitoid lifestyle, most notably in controlling the behaviour of their hosts, but also gene families possibly involved in defense mechanisms against the host’s immune response.

To complete their life cycle, some parasites must influence the behaviour of their hosts. This phenomenon is a notable illustration of the “extended phenotype”, wherein the genes of one organism manifest phenotypic effects in another organism. We found enriched GO terms associated with norepinephrine transport, neurotransmitter transport, and regulation of neurotransmitter activity in Mermithidae. Their genomes exhibit an expansion of gene families that play crucial roles in regulating neurotransmitter levels, motor activity, synaptic transmission, neurotransmitter:sodium symporter activity, GABA-A receptor activity, synaptic vesicle budding from the presynaptic membrane, and the regulation of synapse structural plasticity. Neurotransmission transport was among the expanded gene families shared across nematomorpha, representing an expansion in the common ancestor of nematomorphs. Previous studies identified dysregulated proteins linked to axon/dendrite and synapse modulation that were consistent across hosts of both Mermithidae and Nematomorpha, indicating that neuronal manipulation likely contributes to the induction of positive hydrotaxis (Herbison *et al*. 2019a). Furthermore, neurotransmitter titres were shown to have changed in crickets infected with nematomorph parasites (Thomas *et al*. 2003). Proteomics analysis of the biochemical changes occurring in the cricket *Nemobius sylvestris* and a grasshopper *Meconema thalassinum* when manipulated by the hairworms *Paragordius tricuspidatus* and *Spinochordodes tellinii* respectively identified parasitoid-produced molecules from the Wnt family, which likely affect the host’s central nervous system development. Differential protein expression in the manipulated insects suggested involvement in neurogenesis, circadian rhythm, and neurotransmitter activities (Biron *et al*. 2006, 2005). All of this confirms a genomic capacity of mermithids and nematomorphs to manipulate neuronal connections of their host, possibly either by altering neurotransmitter release in hosts or by producing neuromodulators/neurotransmitters themselves, in a similar way as described for parasitoid wasps (Gal *et al*. 2005; Libersat *et al*. 2009) and tapeworms (Øverli *et al*. 2001).

A less direct way to control the behaviour of its host may be used by the parasitic mermithid *Thaumamermis zealandica*, either instead of or in addition to the neural pathways discussed above. The nematode induces an increase in haemolymph osmolality in parasitised *Bellorchestis quoyana* (formerly *Talorchestia*) comparing to unparasitized amphipods (Williams *et al*. 2004). This alteration may trigger a water-seeking behaviour in the host, providing a potential explanation for their observed tendencies to dig deeper into water saturated sand that the nematode requires to emerge from the host. Our analysis suggests that the last common ancestor of mermithid genomes had expanded gene families associated with potassium ion transmembrane transport and iron ion transport. However, further studies are required to validate the connection between observed patterns of mermithid genomes and manipulation of haemolymph parameters in their hosts. Most descriptions of parasite-induced changes in host behavior rely on observational reports, with limited experimentally confirmed examples of parasite genes directly causing these changes van Houte *et al*. (2013); Biron and Loxdale (2013).

The expansion of gene families associated with various functions such as response to toxic substances, norepinephrine transport, DNA integration, proteolysis, DNA recombination, amino-acid betaine transport, transposition, RNA-mediated mechanisms, and neurotransmitter:sodium symporter activity was observed in both Mermithidae and Nematomorpha, indicating their vital roles in the biology or ecology of both groups of worm parasitoids. Furthermore, this suggests parallel evolution, driven by similar ecological pressures or selective forces, leading to convergent adaptations in phylogenetically distant lineages. These gene families likely contribute to key aspects of the parasitoid lifestyle, such as host immune evasion and manipulation of host physiology Adamo and Webster (2013); Quicke *et al*. (2023); Del Giudice (2019). Parasitoids frequently encounter various host defenses, including the production of toxic substances as part of host immune responses, thus gene families involved in detoxification and responses to such toxins are crucial for parasitic worms to withstand and overcome these challenges. For instance, studies have highlighted the significance of detoxification enzymes like glutathione S-transferases in facilitating the metabolic adjustments required for the endoparasitoid wasp *Pteromalus puparum* to adapt to papilionid butterfly pupae (Xu *et al*. 2020).

We also observed some differences in genomes between Mermithidae and Nematomorpha. For example, both mermithids and nematomorphs accumulate resources during the infective juvenile stage and cease feeding as adults, indicating a shared life history strategy. However, while nematomorphs exhibit enrichment of gene families in response to starvation, indicating potential adaptations to periods of nutrient scarcity, such enrichment is not observed in mermithids. This difference might suggest divergent metabolic strategies between these two groups, warranting further investigation to elucidate the underlying mechanisms and ecological implications. The sensorium in Nematomorpha is practically unknown and the question of how hairworms find mates for internal insemination has never been satisfactorily answered Sokolova *et al*. (2022). Our discovery of expanded gene families enriched in GO terms associated with sensory perception of sound (vibrations) offers an interesting avenue to explore. Mermithids, on the other hand, are fully equipped with a variety of sensory structures, including photoreceptors, although there is limited understanding regarding the expression of phototaxis in different mermithid species. We found expanded gene family related to visual perception in mermithids which might mirror the host-seeking behavior and presence of photosensitive structures in *Mermis nigrescens* Mohamed *et al*. (2007), while information regarding *Romanomermis* remains elusive.

### Transposable elements

TE content displays a wide variation among the studied species, and the underlying reasons for this diversity remain unclear. Particularly, *Nasonia* parasitoid wasp exhibits notably high TE content, yet the factors driving this phenomenon remain elusive Wurm and Keller (2010). The variability of TE landscapes within and across insect orders suggests a complex interplay influenced by factors beyond phylogenetic relatedness, possibly including horizontal transfer. While the impact of insect traits on TE landscapes remains uncertain, recent discoveries suggest TEs play crucial roles in insect adaptations, aging, and antiviral immunity. Thus, TEs are increasingly recognized as essential symbiotic elements in insects, with potentially both harmful and beneficial effects depending on the context Gilbert *et al*. (2021). Therefore, our analysis focuses on determining TE content in the species available to us, aiming to establish foundational data for further investigations.

There are differences in the copy number of TE families between different studied genomes as shown in the (Figure 5). The mermithid genomes have varying completeness with *R. culicivo-rax* and *M. nigrescens* possesing much better contiguity than the genome of *R. iyengari*. Detection and annotation of TEs is sensitive to the quality and contiguity of the assembly. The draft assembly of *Nectonema munidae* has a relatively high contiguity 213 Mbp genome, and an N50 of 716.6 Kbp and that of *Romanomermis iyengari* shows a 336 Mbp genome and an N50 of 152 Kbp, still needing improvement in terms of contiguity and could be the source of inflated TE amounts. Nevertheless, the detection of the TE superfamily composition is not impeded.

Understanding the conserved TE superfamilies within a lineage sheds light on potentially active TEs within the respective group. From our analysis it can be derived that the superfamilies of *hAT, CACTA, Mutator, Transib, Mariner, Helitron* and *Gypsy* persisted successfully among all groups while *Onchocerca volvulus* and *Brugia malayi* succeeded in eliminating whole superfamilies like *PiggyBac, ERV, Jockey* and SINES and downregulating the TE activity overall. Apart from *Nectonema munidae*, it seems that the amounts and activities of TEs are decreasing proportionately in the Nematomorphs. On the contrary they increase in Mermithids, and while generally suppressed in activity and low in amount in the comparative in-group of *Brugia malayi, Onchocerca volvulus* and *Plectus sambesii*, TEs also exhibit a recent burst in activity, most strikingly in *Plectus sambesii. Mermis nigrescens* is the only species where *DIRS* elements were detected.

## Conclusions

Whereas comparative analysis of parasitoid genome sequences alone cannot fully explain parasitoid biology and interactions with hosts, our findings offer foundational insights into the genomic factors driving parasitoid worm adaptation, laying the groundwork and allowing future research to focus on particular gene families. Further, the identification and characterization of transposable element superfamilies within nematomorph and mermithid genomes provide a valuable starting point for a more in-depth exploration of their genomic impact. The significance of studies of gene family evolution and identification of particular gene families associated with specific functions extends beyond mere theoretical insights. They may hold practical value for addressing real-world challenges, particularly in the realm of vector control strategies. Both mermithids and nematomorphs have strong negative effects on the population density of their hosts, in addition to the manipulation of their host’s dispersal behavior discussed above. Not covered in this particular study, but no less important is the ability of mermithids to affect host reproduction success by sterilization, inducing changes in secondary sex characters (intersexes) and mating behavior Vance (1996); Ya’cob *et al*. (2021); Muñoz-Muñoz *et al*. (2016); Rowell (2000). Specifically, uncovering the mechanisms of parasitoid-induced changes in dispersal behavior and reproductive development has a promising potential for applying the knowledge obtained in developing targeted and environmentally friendly methods for controlling insect pests. By leveraging the understanding of molecular factors involved in parasitoid-host interactions, we may uncover innovative approaches to, for example, curbing the transmission of malaria, offering a sustainable and effective means of combating a global health threat.

## Data availability

All data generated in this study are available at https://figshare.com/account/home#/projects/195566.

## Acknowledgments

We would like to thank Professor Dr. Edward Platzer at the University of California Riverside for providing us the culture of *Romanomermis iyengari*.

